# IsoSeq transcriptome assembly of C_3_ panicoid grasses provides tools to study evolutionary change in the Panicoideae

**DOI:** 10.1101/689356

**Authors:** Daniel S. Carvalho, James C. Schnable

## Abstract

The number of plant species with genomic and transcriptomic data has been increasing rapidly. The grasses – Poaceae – have been well represented among species with published reference genomes. However, as a result the genomes of wild grasses are less frequently targeted by sequencing efforts. Sequence data from wild relatives of crop species in the grasses can aid the study of domestication, gene discovery for breeding and crop improvement, and improve our understanding of the evolution of C_4_ photosynthesis. Here we used long read sequencing technology to characterize the transcriptomes of three C_3_ panicoid grass species: *Dichanthelium oligosanthes, Chasmanthium laxum*, and *Hymenachne amplexicaulis*. Based on alignments to the sorghum genome we estimate that assembled consensus transcripts from each species capture between 54.2 and 65.7% of the conserved syntenic gene space in grasses. Genes co-opted into C_4_ were also well represented in this dataset, despite concerns that, because these genes might play roles unrelated to photosynthesis in the target species, they would be expressed at low levels and missed by transcript-based sequencing. A combined analysis using syntenic orthologous genes from grasses with published reference genomes and consensus long read sequences from these wild species was consistent with previously published phylogenies. It is hoped that this data, targeting under represented classes of species within the PACMAD grasses – wild species and species utilizing C_3_ photosynthesis – will aid in futurue studies of domestication and C_4_ evolution by decreasing the evolutionary distance between C_4_ and C_3_ species within this clade, enabling more accurate comparisons associated with evolution of the C_4_ pathway.

## Introduction

The pace of plant genome sequencing has accelerated in recent years. However despite decreases in sequencing costs and improvements in genome assembly quality, species selected for whole genome sequencing often meet one or more of the following criteria: A) agricultural importance, B) status as a genetic model system or C) ecological importance. Sequence data from species which lack direct economic, ecological, or genetic model importance can enable comparative analyses to address biological questions in crops and model species (Michael and Jackson, 2013; Ellegren, 2014). C_4_ photosynthesis has evolved multiple times in the grasses (Edwards et al., 2011), making it particularly amenable to study through comparative genetic approaches (Wang et al., 2009; Huang et al., 2016). C_4_ photosynthesis requires both substantial biochemical and anatomical changes (Kellogg, 2013). All grasses which utilize the C_4_ pathway below to the PACMAD clade, a group of grass subfamilies and tribes which includes substantial numbers of both C_3_ and C_4_ species (Edwards et al., 2011). Substantial new insights into both the genes involved in producing the biochemical and anotomical changes required for C_4_ photosynthesis, as well as the potential function of individual amino acid residues can be obtained from comparative analysis of individual gene families across species utilizing either C_3_ or C_4_ photosynthesis within the PACMAD clade (Christin et al., 2007, 2015; Moreno-Villena et al., 2017). However, assembling sequence data for a single gene family from a large enough set of species through PCR amplification and individual Sanger sequencing remains a time and labor intensive process.

Many domesticated grasses belong to the PACMAD clade, including such as maize (*Zea mays*), sugar cane (*Saccharum spp*.), sorghum (*Sorghum bicolor*), and foxtail millet (*Setaria italica*). However every domesticated grass in the PACMAD clade with a sequenced genome utilizes one or more variants of the C_4_ photosynthetic pathway (Schnable et al., 2009; Garsmeur et al., 2018; Paterson et al., 2009; Bennetzen et al., 2012). As a result, while published whole genome sequence assemblies exist for at least 14 grasses within the PACMAD clade (Table 1), only one of these (*Dichanthelium oligosanthes*, a wild species) (Studer et al., 2016) utilizes C_3_ photosynthesis. Long-read sequencing can effectively generate sequence for large numbers of full length cDNAs even in species lacking reference genome assemblies (An et al., 2018; Zhang et al., 2019). One concern with utilizing this technology for comparative genetic studies is that the higher error rate, particularly the frequencies of insertion and deletion errors, make data from long read based sequencing of non-model species unsuitable for use in comparative evolutionary analyses (Gonzalez-Garay, 2016). However, we previously found that observed synonymous substitution rates calculated from consensus sequences constructed using PacBio IsoSeq pipeline were not elevated relative to a sister lineage where gene sequences were taken from a sanger-based whole genome assembly, indicating sequence data obtained in this manner may indeed be suitable for comparative evolutionary analyses (Yan et al., 2019).

**Table 1:** Published reference genomes for grass species within the PACMAD clade. Species sharing a common inferred evolutionary origin of C_4_ photosynthesis as reported in (Edwards et al., 2011) are indicated by superscript letters.

| Species | Relevance | $C_3/C_4$ | Genome Publication |
| --- | --- | --- | --- |
| *Dichanthelium oligosanthes* | Wild Species | $C_3$ | Studer et al. (2016) |
| *Eleusine coracana*<sup>a</sup> | Grain Crop | $C_4$ | Hittalmani et al. (2017) |
| *Eragrostis tef*<sup>a</sup> | Grain Crop | $C_4$ | Cannarozzi et al. (2014);<br>VanBuren et al. (2019) |
| *Miscanthus x giganteus*<sup>b</sup> | Biomass Crop | $C_4$ | Swaminathan et al. (2010) |
| *Oropetium thomaeum*<sup>a</sup> | Wild Species | $C_4$ | VanBuren et al. (2015, 2018) |
| *Panicum hallii*<sup>c</sup> | Wild Species | $C_4$ | Lovell et al. (2018) |
| *Panicum miliaceum*<sup>c</sup> | Grain Crop | $C_4$ | Zou et al. (2019) |
| *Panicum virgatum*<sup>c</sup> | Biomass Crop | $C_4$ | Casler et al. (2011) |
| *Pennisetum glaucum*<sup>c</sup> | Grain Crop | $C_4$ | Varshney et al. (2017b) |
| *Saccharum spp.*<sup>b</sup> | Sugar Crop | $C_4$ | Garsmeur et al. (2018) |
| *Setaria italica*<sup>c</sup> | Grain Crop | $C_4$ | Bennetzen et al. (2012) |
| *Setaria viridis*<sup>c</sup> | Genetic Model | $C_4$ | Brutnell et al. (2010) |
| *Sorghum bicolor*<sup>b</sup> | Grain/Biomass/<br>Sugar Crop | $C_4$ | Paterson et al. (2009) |
| *Zea mays*<sup>b</sup> | Grain Crop &<br>Genetic Model | $C_4$ | Schnable et al. (2009) |

Here we report the sequencing and characterization of IsoSeq based transcriptomes for three additional PACMAD grasses, selecting to enable wider scale studies of protein sequence changes associated with the many parallel origins of C_4_ photosynthesis within that clade (Figure 1). These species were specifically selected to augment C_3_/C_4_ comparisons: *Hymenachne amplexicaulis, Chasmanthium laxum*, and *D. oligosanthes*. *H. amplexicaulis* is a member of the grass tribe Paspaleae which contains a mixture of C_3_ and C_4_ species. The Paspaleae are sister to an exclusively C_4_ clade consisting of the two grass tribes Andropogoneae + Arundinelleae which include both maize and sorghum, two species with extensive genomic, genetic, and phenotypic resources. *H. amplexicaulis* is found in moist habitats and thrives under flooded conditions (Kibbler and Bahnisch, 1999). *Chasmanthium laxum* belongs to the grass tribe Chasmanthieae (7 species). The Chasmanthieae all appear to utilize C_3_ photosynthesis (Kellogg, 2015) and are generally placed as early diverging lineage within the Panicoideae, the grass sub-family containing maize, sorghum, sugar cane, miscanthus, switchgrass, foxtail millet, and proso millet (Edwards et al., 2011). *C. laxum* can occur in a variety of environments such as: woods, meadows and swamps (Yates, 1966). The final species targeted for transcriptome sequences was *Dichanthelium oligosanthes*. *D. oligosanthes* is the only PACMAD species exclusively utilizing C_3_ photosynthesis with a published genome sequence to date (Studer et al., 2016). It is a member of the grass tribe Paniceae, a group which also includes foxtail millet, proso millet, and switchgrass, but is an outgroup to the MPC C_4_ subclade of exclusively C_4_-utilizing species within that tribe (Giussani et al., 2001; Edwards et al., 2011; Washburn et al., 2015, 2017). As the published *D. oligosanthes* reference genome was constructed utilizing short read sequencing, the inclusion of *D. oligosanthes* provided an opportunity to improve the proportion of genes with full length sequences from this lineage available for comparative analyses. *D. oligosanthes* is present in small glades on the edge of woods (A.J. Studer, personal communication, April 08, 2019). The placement of *C. laxum* as an outgroup to other panicoid grasses with sequenced reference genomes and *D. oligosanthes* as sister to other members of the Paniceae with sequences reference genomes were recovered in a preliminary analysis of our long read dataset. Support of the placement of *H. amplexicaulis* as a sister group to Andropogoneae (sorghum and maize) was strong but not unambiguous.

**Figure 1:**
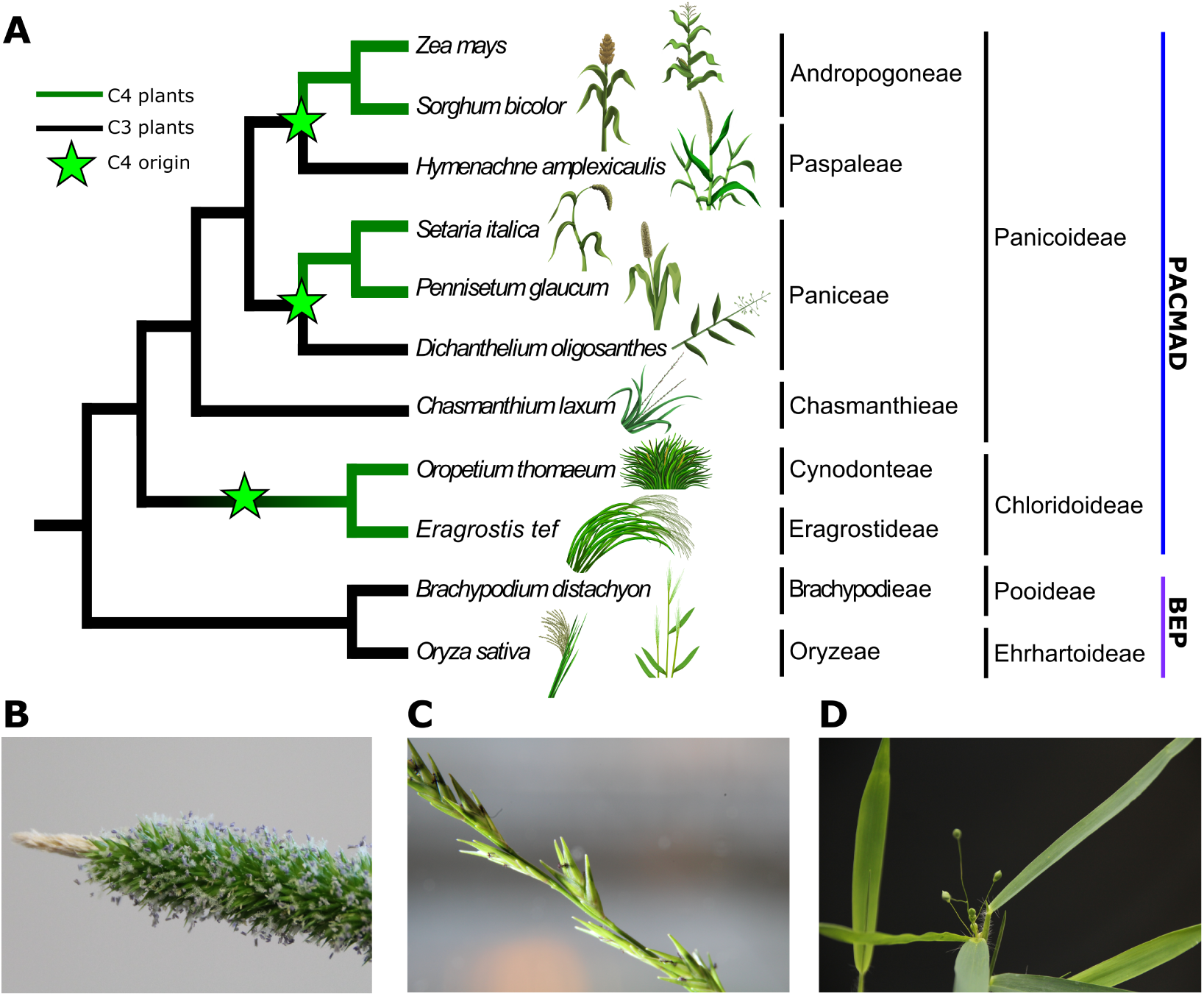
A) Current literature consensus phylogeny of the relationships between the grass species studied here. Lineages in green utilize C_4_ photosynthesis, while lineages in black utilize C_3_ photosynthesis. The green stars indicate apparent independent origins of C_4_ photosynthesis. B) Inflorescence of *H. apmlexicaulis*. C) Inflorescence of *C. laxum*. D) Inflorescence of *D. oligosanthes*.

## Methods

### Plant material, RNA extraction, and sequencing

For all three species, young leaf tissue was harvested from mature plants growing in the greenhouses of the University of Nebraska’s Beadle Center, 40.8190, 96.6932, on October 05 2017. Young leaves were harvested from a *C. laxum* plant germinated from seed collected with accession Kellogg 1268 in Corkwood Conservation Area, just outside of Neelyvile, MO, USA. Full details of this collection are published on Tropicos: https://www.tropicos.org/Specimen/100877982. Leaf tissue from *D. oligosanthes* was harvested from a plant descended from Kellogg 1175, which was collected in Shaw Nature Reserve, west of St. Louis, MO, USA. Full details of this collection are published on Tropicos: http://www.tropicos.org/Specimen/100315254. The specific *D. oligosanthes* plant used as a tissue donor had experienced at least three generations of selfing relative to the originally collected plant. This selfing occurred via an independent lineage from the F2 plant derived from the same collection which was used to generate the DNA for the *D. oligosanthes* reference genome Studer et al. (2016). Young leaves were harvested from *H. amplexicaulis* which had been clonally propagated from collection PH2016. PH2016 was originally collected by Pu Huang in Myakka River state park in Florida, USA on March 22nd, 2016. A clone of this same accession, grown in the same greenhouse, is deposited at the University of Nebraska-Lincoln Herbarium with index number NEB-328848.

Tissue samples were ground in liquid N_2_ and then approximately 200mg of powdered tissue was added to 2 µL of TriPure isolation reagent (Roche Life Science, catalog number #11667157001). The RNA samples mixed with TriPure were then separated using chloroform, precipitated using isopropanol, and RNA pellets were washed using 75% ethanol. The samples were air-dried and diluted in RNAsecure (Ambion). Total RNA concentration was measured using a NanoDrop 1000 spectrophotometer and the integrity was assessed based on electrophoresis on a 1% agarose gel. 10 µL of total RNA for each species was shipped to the Duke Center for Genomic and Computational Biology (GCB), Duke University, USA. Concentrations at the time of shipment ranging from 226.07 to 1,374 ng/µL. One IsoSeq library was constructed per species and each library was sequenced using a single SMRT cell on a PacBio Sequel.

### Consensus reads and transcriptome assembly

Two separate sequence datasets were produced per library: full length (FL) transcripts and non-full length (NFL) transcripts. A given transcript was considered FL if the sequence read contained both 5’ and 3’ adapters as well as poly-A tail and are not redundant to other transcripts. The transcripts lacking the poly-A tail or one of the adapters are instead included in the non-full length dataset. Sequence reads from both files were used to assemble consensus transcriptomes using the software pbtranscript to cluster redundant sequences, part of the SMRT pipe package (version 5.1) with default parameters (https://www.pacb.com/wp-content/uploads/SMRT_Tools_Reference_Guide_v600.pdf). For each final consensus transcript, the single longest ORF present within that transcript was selected as the CDS sequence for downstream analyses. ORFs were required to include an in frame stop codon but were not required to include an in-frame “ATG” which may result in additional non-translated codons being appended to the 5’ end of the putative CDS, but avoids CDS truncation when the 5’ end of the sequence was not recovered.

### Sequence data from species with published reference genomes

CDS file containing only one primary transcript per gene downloaded from Phytozome 12 (https://phytozome.jgi.doe.gov/pz/portal.html) was used from *Brachypodium distachyon* (Initiative et al., 2010), *Oryza sativa* (rice) (Yu et al., 2005; Kawahara et al., 2013), *Sorghum bicolor* (sorghum) (Paterson et al., 2009) and *Setaria italica* (foxtail millet) (Bennetzen et al., 2012). CDS sequences for version 2 of the *Oropetium thomaeum* (oropetium) genome (GenomeID 51527) (VanBuren et al., 2018), and the draft *Eragrostis tef* genome (GenomeID 50954) (VanBuren et al., 2019) were downloaded from CoGe (Lyons and Freeling, 2008). CDS sequences for the initial release of the *Pennisetum glaucum* (pearl millet) genome where downloaded from GigaDB. (Varshney et al., 2017b,a). CDS sequences for B73_RefGenV4 of the *Zea mays* (maize) reference genome was retrieved from Ensembl (Jiao et al., 2017). In cases where only a complete set of CDS sequences was released for a given species, we arbitrarily selected the longest annotated transcript from a given locus to be the single representative transcript for downstream analyses.

### Putative orthology assignments

CDS sequences obtained from *H. amplexicaulis, C. laxum* and *D. oligosanthes* as described above were compared to the primary CDS sequences of each annotated gene in the sorghum genome using LASTZ version 1.04.00 (Harris, 2007) with the following parameters: –identity=70 –coverage=50 –ambiguous=iupac, –notransition, and –seed=match12. CDS sequences from the three target species were presumed to belong to an orthologous group as a given sorghum gene if the sorghum CDS sequence and target species CDS sequence were reciprocally identified as each others high scoring hit in the LASTZ analysis.

Orthologous relationships between sorghum genes and genes in other species with sequenced reference genomes were inferred based on syntenic orthology. For each combination of sorghum and rice, brachypodium, oropetium, teff, foxtail millet, pearl millet, sorghum, and maize all by all LASTZ comparisons were performed using the same parameters described above. The resulting LASTZ output was employed to identify initial syntenic genomic blocks using QuotaAlign with the parameters –tandemNmax=10, cscore=0.5, –merge and –Dm=20 (Tang et al., 2011). The quota was set to –quota=1:2 for maize and teff, and –quota=1:1 for all other species. Pairwise syntenic block data was merged and polished using the methodology previously described in (Zhang et al., 2017) to obtain the final set of high confidence syntenic ortholog groups employed for all downstream analysis.

Orthology was treated as a transitive property, thus each *H. amplexicaulis, C. laxum* or *D. oligosanthes* gene identified as putatively orthologous to a given sorghum gene based on reciprocal best LASTZ hit analysis, was also considered to be putatively orthologous to syntenic orthologs of that sorghum gene identified in each of the other species described above. The final sets of putatively orthologous gene groups including both sequences from published reference genomes and the long read sequencing described here is provided in Supplemental Material 1.

### Sequence alignment, QC, and phylogenetic analysis

Kalign (v2.04) was used create a multiple sequence alignment from protein sequences obtained by translating CDS sequences from all genes in a give putatively orthologous gene group. This gapped protein alignment was in turn employed to create a codon-level DNA alignment of the original CDS sequences. GBlocks version 0.91 was run with default parameters to identify high quality portions of the sequence alignment and remove those portions of the alignment not meeting specified quality thresholds (Talavera and Castresana, 2007). Alignments including only those portions passing GBlocks filering were then used as input for RAxML version 8, using the GTRGAMMA model and with a clade of rice and brachypodium specified as an outgroup, to obtain a phylogenetic tree for each group of putatively orthologous genes (Stamatakis, 2014). When RAxML was unable to construct a phylogeny in which rice and brachypodium formed monophyletic clade sister to other other taxa the trees were omitted from downstream visualization. To plot all phylogenies, we used Densitree, part of the BEAST2 package, was used to create combined blots of large numbers of trees (Bouckaert et al., 2014). For visualization purposes only, all branches were treated as having equal length in order to improve the ease of visually comparing differences in topology.

As a result of the separate whole genome duplications in the maize and teff lineages, in many cases gene and species trees would contain different numbers of leaf nodes. For gene groups where maize and teff had each fractionated back to single copy status, only a single alignment file was created. If fractionation had already occurred in one lineage, but not the other, two separate alignments were created, each sampling one of the two co-orthologous gene copies from the species with a retained whole genome duplication derived gene pair. When fractionation had not occurred in either lineage, four total alignments were generated per gene group, capturing all possible pairwise combinations of the two teff gene copies and two maize gene copies.

## Results and Discussion

The number of raw reads generated per species was largely consistent and ranged from 708,681 to 734,932 (Table 2). After clustering both full length and non-full length transcripts to obtain a set of polished consensus transcripts, the number of sequences per species dropped to 164,640 to 193,422 (Table 2). The average length of consensus sequences ranged from 925 bp to 1,438 kb (Figure S1). The number of consensus transcripts significantly exceeded the expected number of expressed genes, however, this is consistent with other reference genome-free IsoSeq analyses (Li et al., 2017; Kuang et al., 2019; Yan et al., 2019). Inflated numbers of consensus transcripts can result from sequencing of multiple alternatively spliced isoforms of the same gene, sequencing of incompletely processed mRNA molecules (Martin et al., 2014), high sequence error rates preventing multiple sequences from the same transcript being collapsed into a consensus, divergent haplotypes of the same locus present in our clonally propagated, wild collected, or partially inbred starting material, or contamination of the original samples with mRNA from non-target organisms.

**Table 2:** Summary statistics for raw and processed long read sequence data generated from each of the three target species.

| Species | Total reads | Raw data | CCS reads | FL reads | Average FL length | Consensus transcripts | Average consensus transcript length |
| --- | --- | --- | --- | --- | --- | --- | --- |
| *Hymenachne amplexicaulis* | 734,932 reads | 5.8 GB | 732,158 reads | 284,027 reads | 963 bp | 193,422 | 925 bp |
| *Dichanthelium oligosanthes* | 708,681 reads | 10.1 GB | 701,802 reads | 380,381 reads | 1,460 bp | 190,632 | 1,438 bp |
| *Chasmanthium laxum* | 729,710 reads | 12.5 GB | 649,149 reads | 306,566 reads | 1,294 bp | 164,640 | 1,236 bp |

Alignment of final consensus reads to the sorghum reference genome was employed to estimate coverage of the shared grass gene space for data collected from each of our target species, as well as to assist in further collapsing multiple redundant sequences originating from alternative splicing, incomplete processing, or divergent haplotypes of transcripts originating from a single genetic locus. In all three cases that majority of consensus transcripts could be aligned to known genes in the sorghum genome, with an average of between 9.3 and 12.1 consensus transcripts aligning to each sorghum gene represented in the transcriptome data (Table 3). Each of these three target species is predicted to be diploid based on either flow cytometry based estimates of genome size and/or imaging of chromosomes, thus a maximum of two transcripts per locus can be explained by divergent haplotypes. The high number of consensus sequences aligned per represented sorghum locus suggests that a large proportion of the overall inflation in consensus transcript number from this dataset may result from alternative splice isoforms or sequencing of incompletely processed mRNA molecules. It should also be noted that this analysis will confound lineage specific gene duplications with divergent haplotypes and splice isoforms, however this bias will be consistent across all three species.

**Table 3:** Alignment rates of consensus transcripts generated from each of the three target species to the sorghum gene space.

| Species | Sorghum gene space coverage | Sorghum syntenic gene space coverage | Transcript alignment rate |
| --- | --- | --- | --- |
| *H. amplexicaulis* | 11,485 genes/34,211 genes (33.5%) | 6,402 transcripts/11,800 genes (54.2%) | 115,361 transcripts/193,422 transcripts (59.6%) |
| *C. laxum* | 13,446 genes/34,211 genes (39.3%) | 7,418 transcripts/11,800 genes (62.8%) | 125,357 transcripts/164,640 transcripts (76.1%) |
| *D. oligosanthes* | 14,159 genes/34,211 genes (41.3%) | 7,760 transcripts/11,800 genes (65.7%) | 171,465 transcripts/190,632 transcripts (89.9%) |

For each sorghum gene which aligned to two or more consensus transcripts from the same target species, a single representative transcript was selected for further downstream analysis (See Methods). Between 11,485 and 14,159 sorghum genes had a corresponding representative transcript in a given target species (Table 3). Here we were using only single library was constructed per species, rather than multiple libraries constructed using different size fractions, the use of RNA from a single tissue rather than pooled RNA from multiple tissue types, and were conducting comparisons between more distantly related species. However, the total proportion of sorghum genes represented in each transcriptome dataset was not substantially lower than the 14,401 *T. dactyloides*-maize gene pairs identified in a previous study which implemented all of these best practices (Yan et al., 2019). This may in part be explained by both sequencing and library preparation improvements between the RSII and Sequel iterations of this sequencing technology.

Manual curation was used to access the coverage and quality of sequences retrieved from these three C_3_ photosynthesis-utilizing PACMAD species for five genes known to be involved in C_4_ photosynthesis: PPDK, PEPC, NADP-MDH, NAD-ME and DCT2 in C_4_ photosynthesis-utilizing PACMAD species. In four cases, the representative transcript identified from each of the three target species spanned every annotated codon in sorghum. The one exception was PPDK where the representative transcript identified for *H. amplexicaulis* lacked the first annotated exon of the annotated gene model in sorghum (Figure 2). Multiple isoforms of the PPDK gene have been described in both maize and sorghum, with the shorter isoform, lacking the same exon absent in *H. amplexicaulis* (Sheen, 1991; Wang et al., 2009). This shorter isoform lacks the chloroplast transit peptide and encodes cytosolic PPDK protein not thought to be associated with C4 photosynthesis (Glackin and Grula, 1990; Sheen, 1991; Wang et al., 2009).

**Figure 2:**
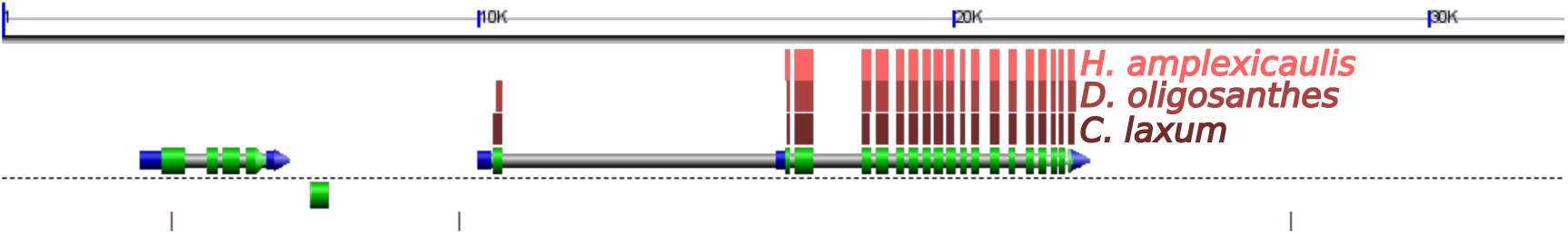
A GEvo panel showing transcript coverage of the C_4_ PPDK gene in *S. bicolor* Sobic.009G132900 in each of the three species texted. Red-brown boxes represent regions of similar sequence identified by BLASTN between the sorghum genome and consensus transcript sequences retrieved from *H. amplexicaulis, D. oligosanthes, C. laxum* (from top most to bottom most). The bottom track indicates the annotated gene structure, with intronic sequence indicated in gray and exonic sequence indicated in either blue (5’ or 3’ untranslated regions) or green (coding sequence). Top y-axis indicates scale of the displayed genomic region in kilobases. (Lyons and Freeling, 2008).

Phylogenetic consistency was assessed using a small subset of genes with high confidence syntenic orthologs identified in species with published reference genomes and representative transcripts identified in each of the three target species. A total of 11,800 genes were identified at syntenic orthologous locations across the genomes of rice, brachypodium, teff, oropetium, pearl millet, foxtail millet, sorghum, and maize. Of these in 2,774 cases no representative transcripts were retrieved from *C. laxum, D. oligosanthes*, or *H. amplexicaulis*. These cases likely represent conserved genes that are not expressed in developing photosynthetic tissue. In 1,611 cases, a representative transcript was identified in only one of the three target species, and in 2,276 cases, representative transcripts were identified in two of the three target species. In the remaining 5,139 cases representative transcripts were retrieved for all three target species. The complete lists of each of these sets of conserved syntenic genes and corresponding transcripts from 0, 1, 2, or 3 of the target species is provided as part of Supplemental Material 1.

One potential concern is using transcriptome data from species utilizing C_3_ photosynthesis to provide sequence data for comparative genetic and evolutionary analyses of C_4_ is that enzymes involved in the C_4_ cycle will likely different functions unrelated to photosynthesis in C_3_ plants (Aubry et al., 2011), and therefore may not be expressed in photosynthetic tissue and hence be missing from from datasets derived from sequencing cDNAs. Of 31 core C_4_ genes enumerated in (Huang et al., 2016), 20 were part of the set of 11,800 sorghum genes with conserved syntenic orthologs identified in each of the tested grass species with a published reference genome. Hence, these genes are almost certainly present within the genomes of *C. laxum, D. oligosanthes*, and *H. amplexicaulis* as well, whether or not they were expressed to sufficient levels to be detected in this analysis. Of these 20 syntenically-conserved C_4_ related genes, sequence data was obtained from all three target C_3_ utilizing panicoid species in 16 cases. In the remaining four cases – DCT4c, GLR, NADP-ME and SCL – no putatively orthologous transcript was identified in any of the three species. There were no cases where a syntenically conserved gene linked to C_4_ photosynthesis was detected in some, but not all, of the three C_3_ utilizing species evaluated.

From the list containing a total of 5,139 conserved orthologous gene groups present in all species 231 were discarded for one of several reasons, listed from most common to least common. 1) In 113 cases the CDS sequence for the *O. thomaeum* genome included one or more in-frame stop codons. 2) In 61 cases in at least one species represented by isoseq data no stop codon was present in any of the 6 possible open reading frames, indicating either a sequencing error or incomplete 3 prime coverage. 3) In 56 cases a syntenic orthologous gene present in version 2.1 of the *B. distachyon* genome had been removed or renamed in version 3.1 of the *B. distachyon* genome. 4) One *O. thomaeum de novo* predicted gene region was not present in the CDS data.

The remaining set of 4,908 conserved orthologous gene groups were used to generate protein-guided codon multiple sequence alignments (See Methods). A subset of these alignments containing at least 900 nucleotides (300 codons) alignment scored as “high quality” by GBlocks were employed to construct individual gene-level trees (Figure S2). In total 746 trees, representing 275 putatively orthologous gene groups were constructed. Multiple trees resulted from retained duplicate gene pairs resulting from lineage specific whole genome duplications in maize and teff. Each duplication had the potential to create a retained syntenic gene pair which were each co-orthologous to single gene copies in other grass species within the analysis. In order to maintain a consistent number of final nodes, when a retained gene pair was observed in one or both species, multiple sampled trees were generated (see Methods). A modest bias towards over representation of retained – rather than fractionated – genes was observed in the set of genes which were represented in the transcriptome assemblies from all three target species: 37% (1,843/4,908) of maize genes in this set were retained as duplicate pairs vs 30% of all syntenic maize genes, and 100% of teff genes in this set were retained as duplicate pairs vs 91% of all syntenic teff genes. The rice and brachypodium clade represented a known outgroup as these two species belong to the BEP clade which diverged from the PACMAD clade of grasses early in the evolution of this family (Edwards et al., 2011). In 46 cases, RaxML was unable to place the rice-brachypodium clade as an outgroup suggested broader issues with orthology assignment, correct ORF identification, or alignment. These trees were not included in downstream analyses.

Among the 700 remaining gene trees, 304 (43%) produced a single topology consistent with the prior literature on the relationship of these species (Figure 3). The second and third most common topologies were each represented by less than 7% of all calculated trees, 47 and 44 cases respectively. The second and third most common topologies differed from prior published phylogenies regarding the placement *H. amplexicaulis*. In the second most common topology *H. amplexicaulis* was placed sister to all other panicoid grass species other than *C. laxum*. In the third most common topology *H. amplexicaulis* was placed sister to the Paniceae. Parallel analysis was conducted using all 4,908 conserved orthologous gene groups, including many cases with substantially shorter regions of high quality multiple sequence alignment. The pattern of trees recovered were largely consistent with those in (Figure 3). In the “all genes” analysis, the same most common topology was retrieved as in the long alignment only analysis. The second most common topology in the “all genes” analysis corresponds to the most third most common topology in Figure 3, while the third most common in the “all genes” topology places *C. laxum* as sister to the combined Chloridoideae and Panicoideae (Supplemental Figure S3).

**Figure 3:**
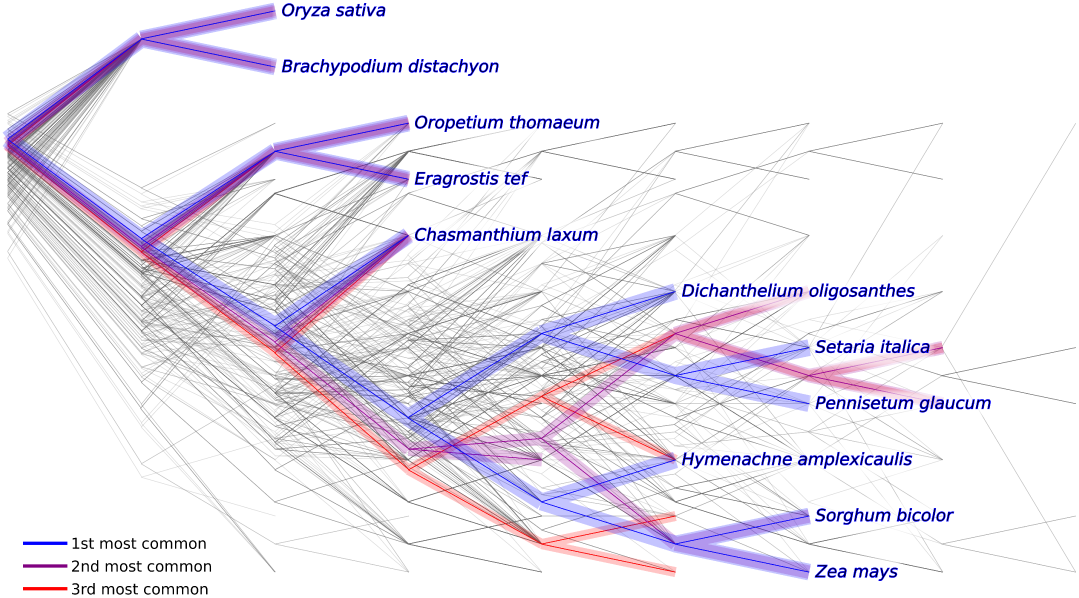
Seven hundred distinct phylogenetic trees calculated from separate multiple sequence alignments of 275 putatively orthologous gene groups with large regions of alignment scored as high quality. Blue indicates the most commonly observed topology (304 trees (43% of the total), purple and red indicate the second (47 trees (6.7%) and third most commonly observed topologies (44 trees (6.2%)), respectively.

## Data availability

Raw sequence data for *C. laxum, H. amplexicaulis* and *D. oligosanthes* have been deposited in the NCBI SRA under accessions numbers: SRR7632721 (*C. laxum*), SRR7632716 (*H. amplexicaulis*) and SRR9603193 (D. oligosanthes). Processed consensus transcript sequences generated for each of these three species have been deposited in Zenodo DOI 10.5281/zenodo.3253206. Syntenic gene sets and putatively orthologous relationships are provided as Supplemental Material 1.

## Author’s contributions

DSC and JSC wrote the paper and designed the experiments, DSC generated and analyzed the transcriptome data. All authors have reviewed and approved the manuscript.

## Acknowledgements

This work was supported by a CNPq Science Without Borders scholarship (214038/2014-9) to DSC. This material is based upon work supported by the National Science Foundation under Grant No. MCB-183830 to JCS. This project was supported by the Agriculture and Food Research Initiative Grant number 2016-67013-2461 from the USDA National Institute of Food and Agriculture to JCS. In addition, portions of this project were supported by Robert B. Daugherty Water for Food Institute research support award to JCS. The authors would like to thank Lindsay Erndwein who drew the illustrations of various grass species presented in Figure 1A.

## Supplemental figures

**Figure S1:**
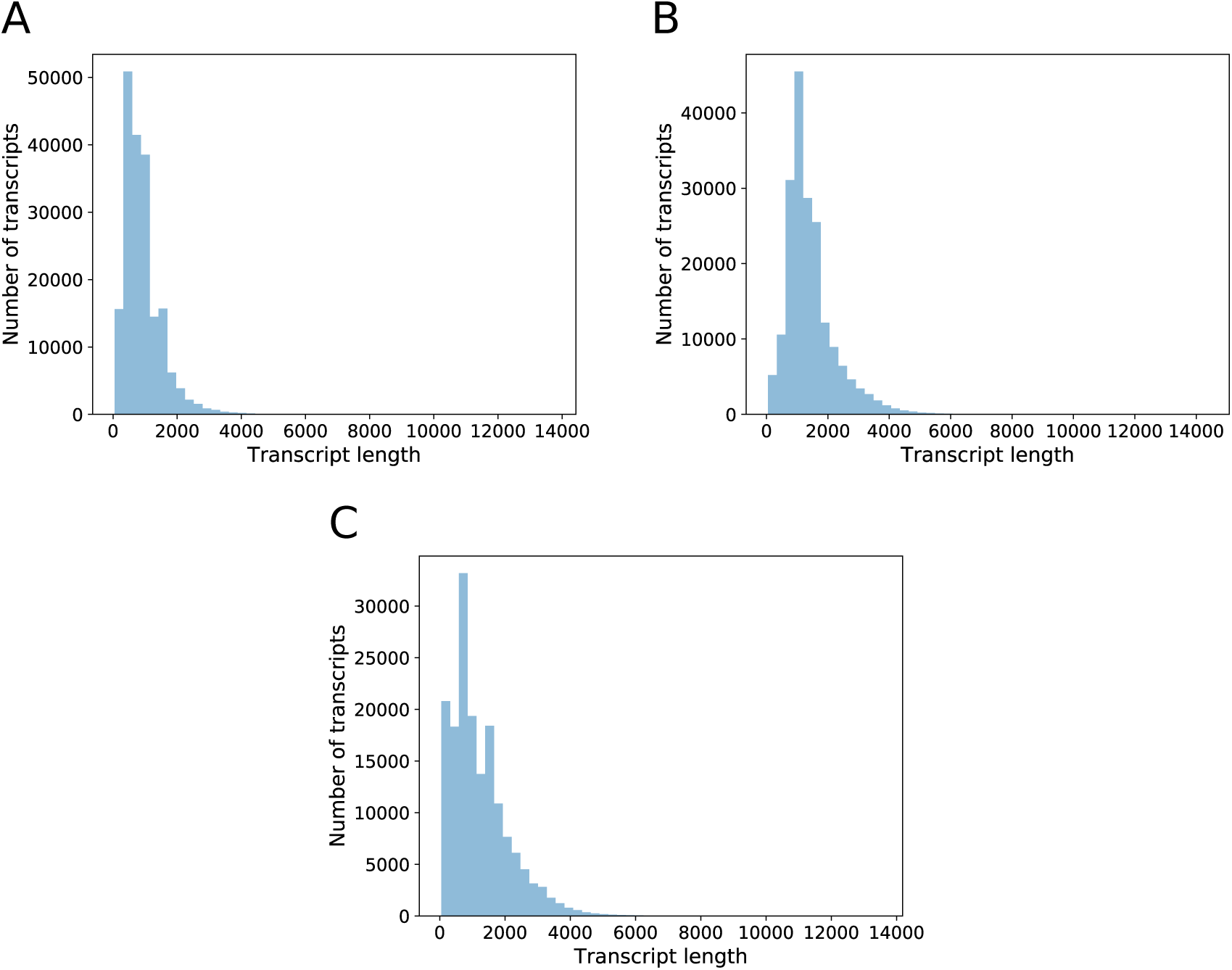
Distribution of lengths for polished transcript sequences: A) *H. amplexicaulis*, B) *D. oligosanthes*, C) *C. laxum*.

**Figure S2:**
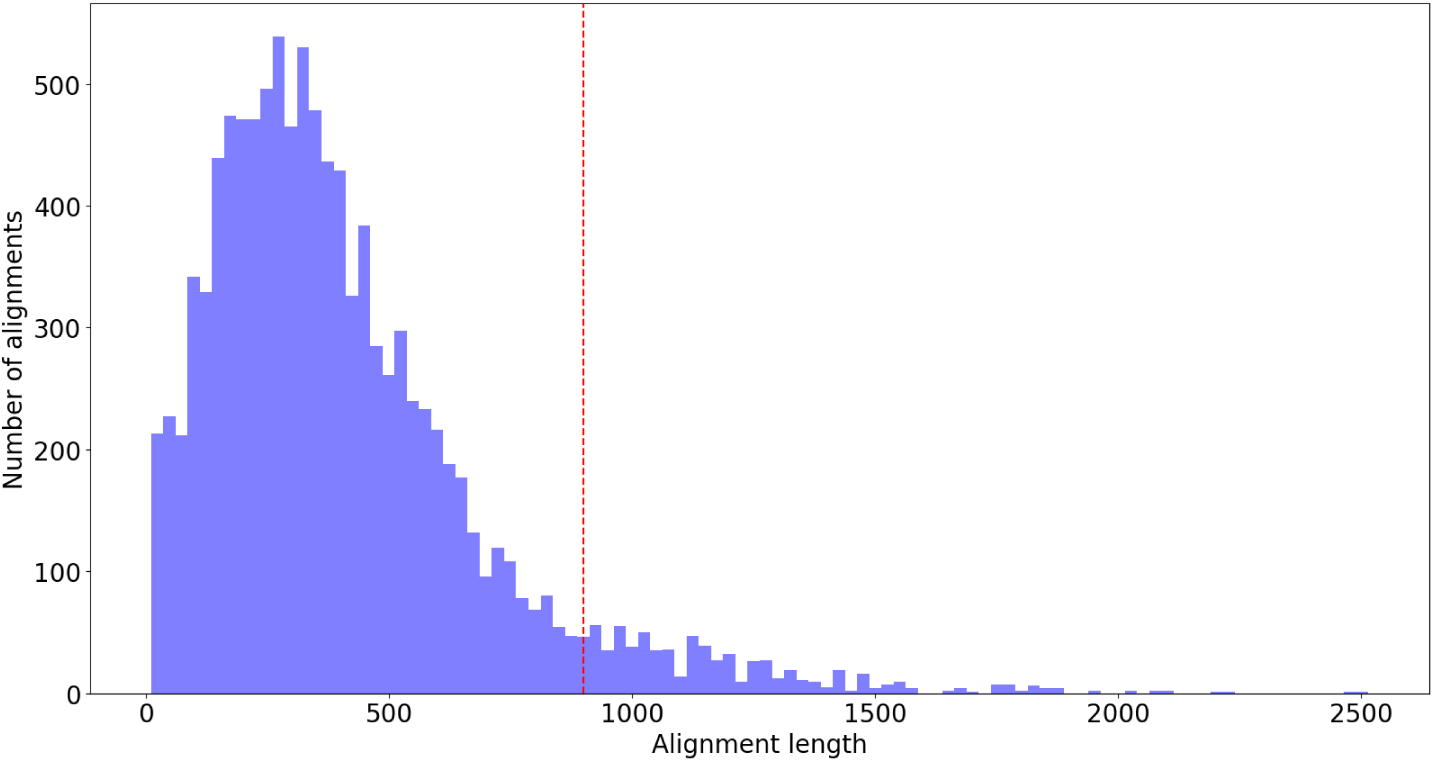
Distribution of alignment lengths after GBlocks cleaning. Dashed line represents the threshold of 900 nucleotides long sequences employed for Figure 3. Sequences represented on the right side of the histogram were analyzed.

**Figure S3:**
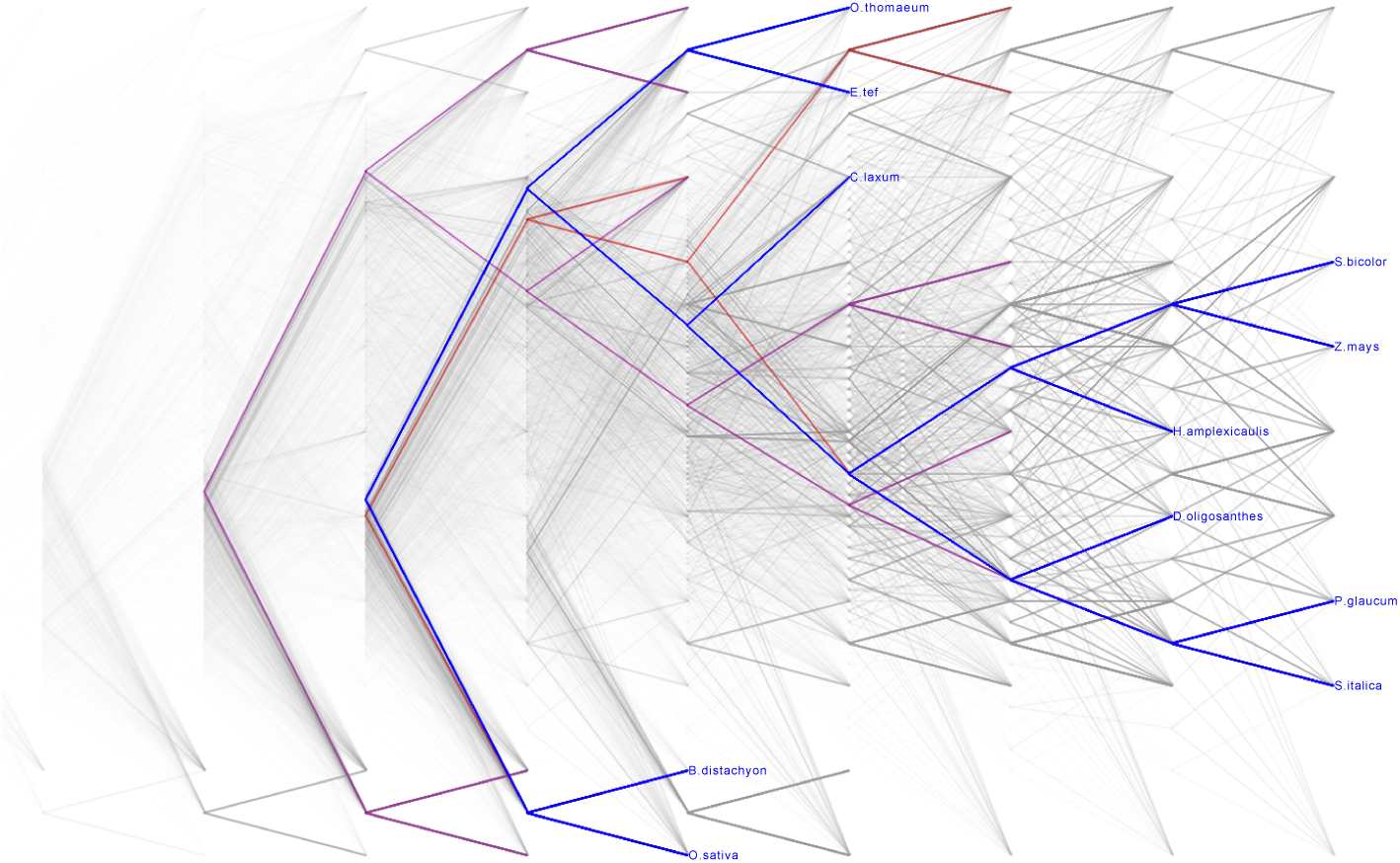
Plot of 10,876 phylogenetic trees. The blue branches represent the most common topology, purple and red branches represent second and third most common topologies, respectively. Figure generated using Densitree.

